# Machine learning application to predict binding affinity between peptide containing non-canonical amino acids and HLA0201

**DOI:** 10.1101/2024.11.19.624425

**Authors:** Shan Jiang, Zhaoqian Su, Nathaniel Bloodworth, Yunchao Liu, Cristina Martina, David G. Harrison, Jens Meiler

**Author notes:** Equal contribution (Shan Jiang, Zhaoqian Su).

## Abstract

Class 1 major histocompatibility complexes (MHC-I), encoded by the highly polymorphic HLA-A, HLA-B, and HLA-C genes in humans, are expressed on all nucleated cells. Both self and foreign proteins are processed to peptides of 8 to 10 amino acids, loaded into MCH-1 within the endoplasmic reticulum and then presented on the cell surface. Foreign peptides presented in this fashion activate CD8+ T cells and their immunogenicity correlates with their affinity for the MHC-1 binding groove. Thus, predicting antigen binding affinity for MHC-I is a valuable tool for identifying potentially immunogenic antigens. While quite a few predictors for MHC-I binding exist, there are no currently available tools that can predict antigen/MHC-I binding affinity for antigens with explicitly labeled post-translational modifications or unusual/non-canonical amino acids (NCAAs). However, such modifications are increasingly recognized as critical mediators of peptide immunogenicity. In this work, we propose a machine learning application that quantifies the binding affinity of epitopes containing NCAAs to MHC-I and compares its performance with other commonly used regressors. Our model demonstrates robust performance, with 5-fold cross-validation yielding an R^2^ value of 0.477 and a root-mean-square error (RMSE) of 0.735, indicating strong predictive capability for peptides with NCAAs. This work provides a valuable tool for the computational design and optimization of peptides incorporating NCAAs, potentially accelerating the development of novel peptide-based therapeutics with enhanced properties and efficacy.

## Introduction

The class I major histocompatibility complex (MHC-I) enables the adaptive immune response by presenting antigens to patrolling cytotoxic T cells [1,2]. Peptides presented by MHC-I originate in the cytoplasm and are usually length limited, having only 8 to 10 amino acids. This system has evolved principally to enable rapid identification and elimination of viral infected or malignant cells while minimizing the risk of aberrant recognition of self-peptides and consequential autoimmunity [3]. The MHC-I protein products are themselves encoded by the Human Leukocyte Antigen (HLA) genes in humans; both the co-dominantly expressed subtypes (A, B, and C) and the high degree of polymorphism observed in the peptide-binding domain of these genes enable MHC-I to complex with a large repertoire of peptides [3,4]. Post-translational modification of proteins and peptides (resulting in the incorporation of NCAAs) can further broaden the immunogenic landscape of peptides presented by MHC-I. Peptides containing various NCAAs are implicated as immunogens in a variety of diseases [1] including rheumatoid arthritis [5], hypertension (ref PMID 25244096 and PMID 39145457) and cardiometabolic inflammation [6], and cancer [7].

Recent advances in immunotherapy targeting cancer and autoimmune diseases, coupled with advances in data science have incentivized the creation of computational tools that predict peptides likely to bind to MHC-I and induce immune responses [8,9]. These tools encompass a wide range of methodologies to analyze peptide-MHC interactions. Among these are sequence-based approaches like NetMHCPan [10–12] and MHCflurry [13] that utilize amino acid sequences to forecast binding affinities. Additionally, structure-based approaches such as Rosetta FlexPepDock [14–16] employ three-dimensional structural data to provide a detailed understanding of peptide-MHC binding dynamics and conformational stability. The most advanced and effective of these tools leverage machine learning techniques to construct predictive models. These models are trained on extensive datasets comprising antigen-MHC-I pairs and their corresponding binding affinity data. A significant portion of these data are derived from the Immune Epitope Database (IEDB), which provides a comprehensive repository of experimentally validated immune epitopes. These methods are thoroughly benchmarked and reviewed by Zhao et al [8].

Despite notable advances in both sequence- and structure-based epitope binding predictors, there are currently no tools capable of rapidly predicting antigen/MHC-I binding affinity for antigens with post-translational modifications or NCAAs. These modifications are increasingly recognized as critical mediators of peptide immunogenicity. The main scope of this research is to develop a new model that would be able to predict the binding affinity of epitopes containing NCAAs to MHC-I.

Machine learning models have demonstrated superior performance in predicting binding affinity due to their ability to capture complex patterns and interactions within the data. The development and refinement of these models involve rigorous processes including feature generation, model training, and validation. Several popular algorithms are widely used for property prediction in the fields of chemistry and biology, including support vector machines (SVM), artificial neural networks (ANNs), principal component analysis (PCA), and partial least squares (PLS) regression [17,18].

In this work, we develop a simple encoder capable of creating feature vectors from peptides based on chemical structure. We then systematically benchmark several different supervised and unsupervised machine learning models on a filtered, publicly available dataset containing peptides with NCAAs and experimentally determined binding affinities.

## Results

This section details the study results from three perspectives, data preparation, feature generation and model testing and validation. The data preparation subsection will explain the source and structure of the data used, focusing on data exploration and filtration. The feature generation subsection, the key part of this section, will introduce how peptides with NCAAs are encoded. The model testing and validation subsection will evaluate and compare performance metrics such as R^2^ and RMSE across different datasets with five-fold cross-validation using various algorithms.

### Data preparation

The initial dataset, a table with 100,141 rows and 29 columns, was exported from the publicly available Immune Epitope Database (IEDB). Among the 29 columns, five are of particular interest for this study: “Name”, “Qualitative Measurement”, “Quantitative Measurement”, “Response Measured”, and “HLA”. Table 1 lists possible or example values for these five columns. The “Name” column shows the peptide sequence within the binding complex of interest; in the given “Name” column example in the table, GILGFVFTL + OTH(L9), the text between the “+” sign and parentheses indicates the modification method applied to the peptide, and the text within the parentheses lists the amino acids modified by this method. The “HLA” column shows the HLA gene responsible for encoding the MHC binding to the peptide. The “Qualitative Measurement” column has values ranging from strong to weak, representing binding strength. The “Quantitative Measurement” column provides a numerical value obtained from experiments, with the type of measurement explained in the “Response Measured” column.

**Table 1:** Table listing the five columns of most interests, example values, and the number of unique values.

| Column Name | Example Values | Number of Unique Values |
| --- | --- | --- |
| Name | GILGFVFTV + OTH(V9) | 61813 |
| Qualitative Measurement | positive, negative | 5 |
| Quantitative Measurement | 0.1 – 65,000 | 223 |
| HLA | HLA-A*02:01 | 122 |
| Response Measured | half maximal inhibitory concentration (IC50) | 8 |

Figure 1 demonstrates the data preparation process. Starting with the original dataset of 100,141 rows, it was confirmed that each peptide contains at least one NCAA. Since the objective of our research is to predict quantitative binding affinity, each row needed a non-NA value in the “Quantitative Measurement” column. Additionally, to ensure consistency of “HLA” and “Response Measured” across the training and test datasets, the most populated “HLA” and “Response Measured” values, which were HLA-A*02:01 and IC50 with a unit of nM were selected. Finally, a dataset of 166 rows was prepared for further analysis.

**Figure 1:**
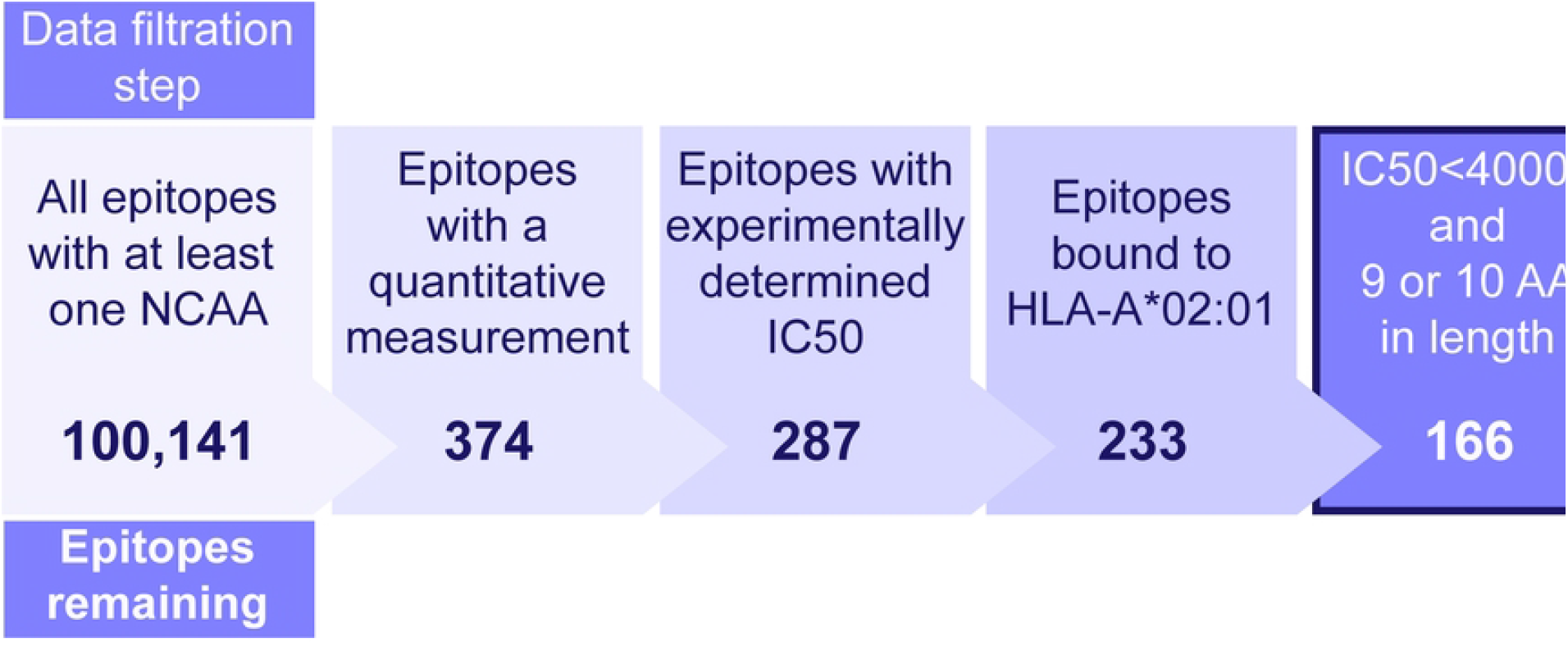
Dataset generation for model training and validation. Epitopes with experimentally determined IC50 values were extracted from the IEDB and filtered as shown to generate the dataset used to generate the model.

### Feature generation

With the sequences of peptides and their quantitative binding values prepared, the next step was to determine how to encode them for machine learning model building. Protein/amino acid encoding involves representing a protein or amino acid with an n-dimensional numerical vector. According to published studies, there are multiple encoding methods, which can be either whole sequence-based or amino acid-based [19]. In the latter approach, each amino acid is first encoded individually, and then the combination of feature vectors from all amino acids in the protein sequence constitutes the encoding of the entire peptide sequence.

Since all HLA species across the dataset used for this study are the same, only the peptides of the binding complex need to be considered for generating the input vector for the next step’s model building. This simplifies the process and makes it more time efficient. Given that the target peptides in this study contain at least one NCAA, which implies potential chemical modifications at the same amino acid position, it is intuitive to use chemistry or structural encoding rather than sequence encoding to retain residue-specific information.

The feature generation process includes four main steps, as illustrated in Figure 2. First, each peptide sequence is tokenized into amino acid tokens. According to summary after step 1, for all 166 rows of data, totally 20 canonical and 28 non-canonical tokens were generated. Figure 3 shows the count of unique tokens across the entire dataset, while Figure 4 illustrates the distribution of tokens at each amino acid position. Token name that contains “_” indicates a NCAA. Second, the structure of each amino acid token, particularly NCAAs, were verified using referenced literature searched from IEDB by “Epitope IRI”, and chemical structures are converted to SMILES strings. Third, feature vectors for each token were generated using RDKit from the SMILES strings obtained in the previous step. According to RDKit, these vectors describe various physicochemical properties such as molecular weight, partial charge, and the number of specific functional groups, resulting in a total of 208 features. Given the size of the prepared dataset, the feature vector dimension is large, and many features are highly correlated, so principal component analysis (PCA) was applied to reduce the dimensionality from 208 to 10. The choice of 10 components was made because they cover 99.75% of the variance of the original feature vector.

**Figure 2:**
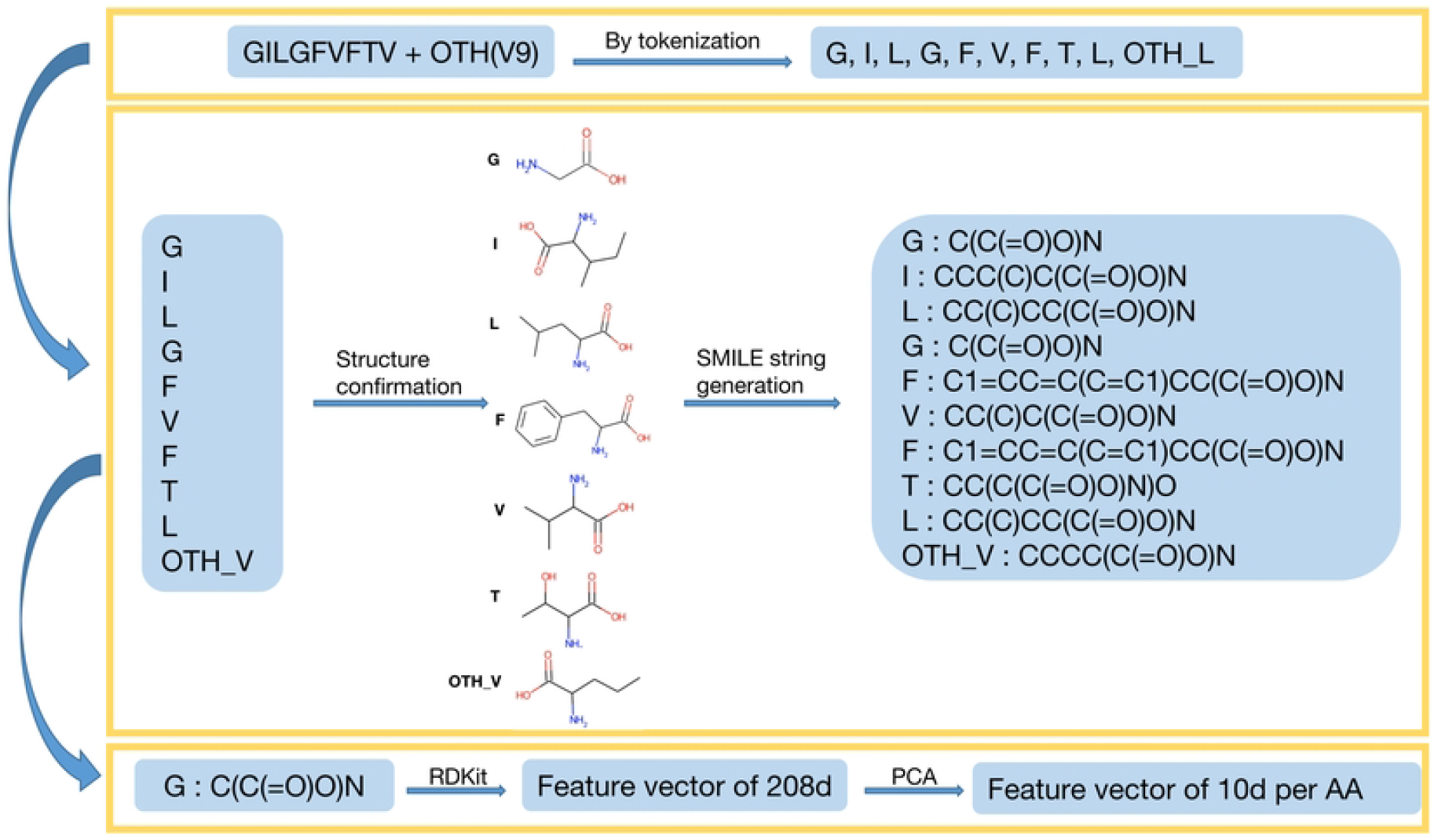
Process for encoding naturally occurring and non-canonical amino acids. Peptides were tokenized by individual amino acid, structures of NCAAs manually confirmed, and SMILE strings for each structural representation generated. These SMILE strings were vectorized using RDKit followed by feature reduction with PCA.

**Figure 3:**
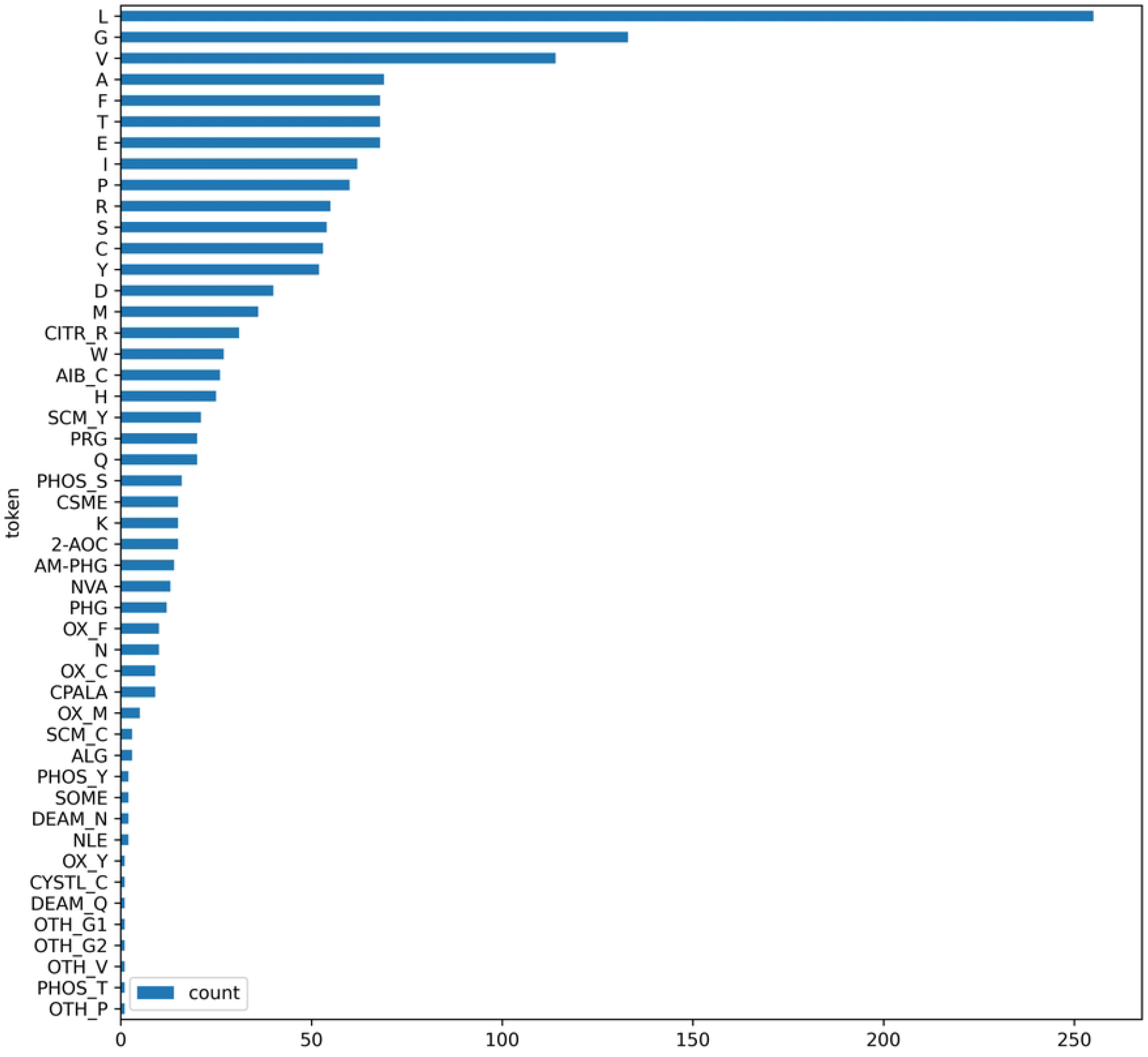
Distribution of canonical and NCAA tokens for every epitope in the dataset.

**Figure 4:**
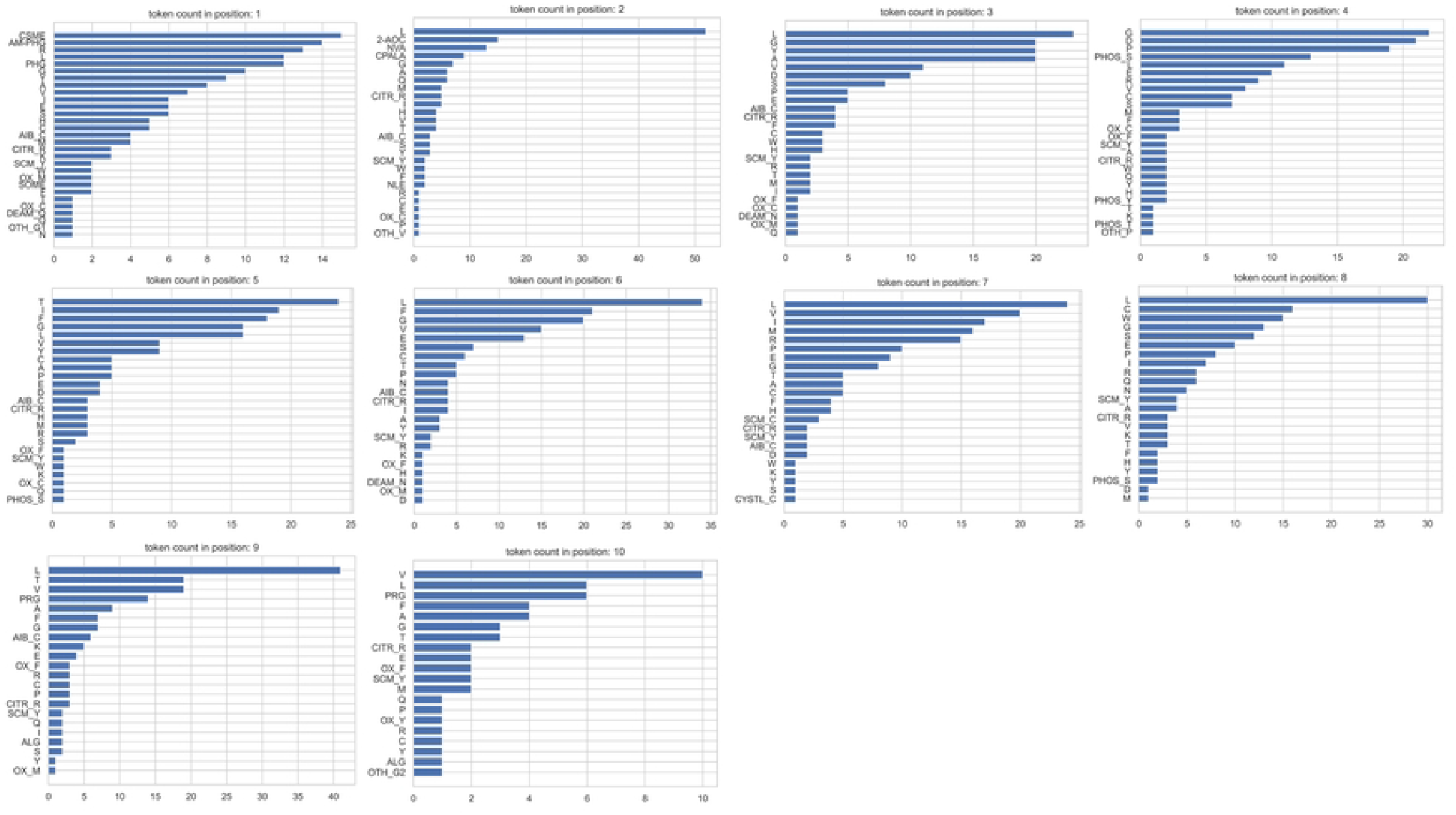
Distribution of canonical and NCAA tokens at each residue position (labeled N to C terminus).

At this point, a map was created with an amino acid token as the key and its corresponding feature vector of size 10 as the value. The final step is to combine the features of all tokens obtained in the first step to generate the feature vector for the entire peptide sequence. With each token’s vector size being 10 and the peptide length being nine or ten, the resulting feature vector dimensions for each peptide sequence would be 90 or 100. To ensure consistency of input data for building a machine learning model, an additional 10 zeros were appended to the feature vectors of peptides with a length of nine.

### Model testing and validation

Figure 5 demonstrates the framework of the model. Each input is a feature vector with 100 dimensions derived from the peptide, and the output is the logarithm of IC50 (nM). During model building, a five-fold cross-validation was applied to the dataset. Root mean square error (RMSE) and R-square (R^2^) were used as evaluation metrics. To compare the training results of Partial Least Squares (PLS) with other commonly used algorithms, an open-source tool named Lazy Predict was applied to the same dataset.

**Figure 5:**
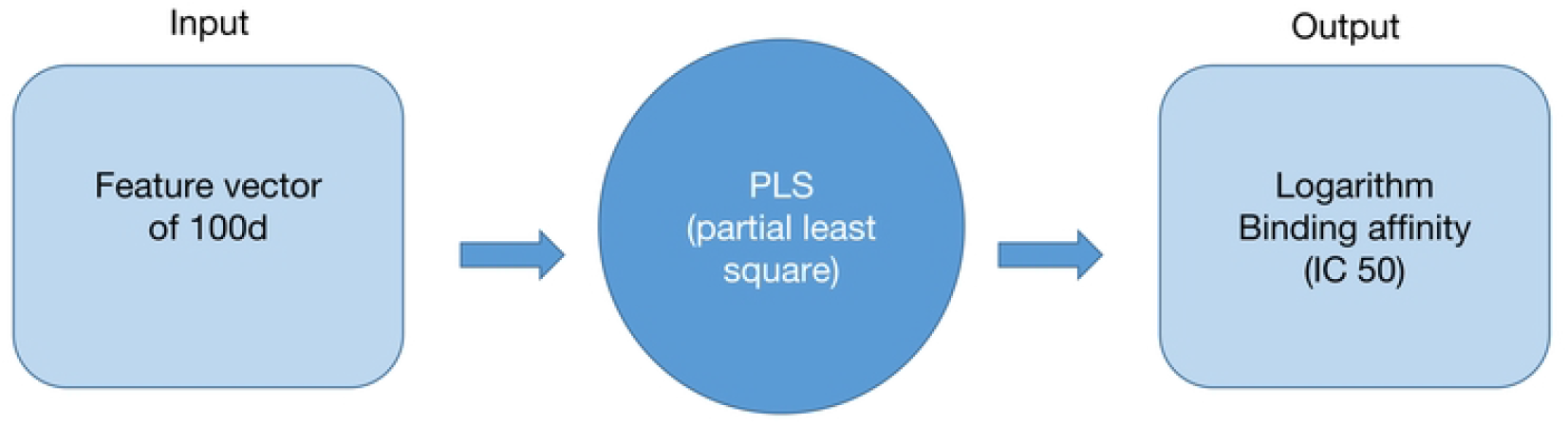
Overview of the predictive model framework.

Three components were selected for building the Partial Least Squares (PLS) model because, among the range of 2 to 10 components tested, using 3 components yielded the best performance in terms of cross-validated R-squared (R^2^) and root mean square error (RMSE) using five-folds. The detailed results of this comparison are listed in Table 2. Figure 6 illustrates the correlation between the original binding affinity and the predicted binding affinity for both the training set and the test set, using PLS from each individual cycle of five-fold cross-validation. The scatter plots reveal a clear correlation between the actual and predicted values, demonstrating the model’s effectiveness despite the relatively small dataset. This strong correlation in both training and test datasets indicates that the model generalizes well and is not overfitted.

**Table 2:** Performance of PLS with different components - cross-validated R^2^ and RMSE.

| Components | Cross-validated $R^2$ | Cross-validated RMSE |
| --- | --- | --- |
| 2 | 0.444 | 0.759 |
| **3** | **0.477** | **0.735** |
| 4 | 0.463 | 0.743 |
| 5 | 0.451 | 0.750 |
| 6 | 0.425 | 0.766 |
| 7 | 0.395 | 0.786 |
| 8 | 0.355 | 0.810 |
| 9 | 0.325 | 0.827 |
| 10 | 0.283 | 0.852 |

**Figure 6:**
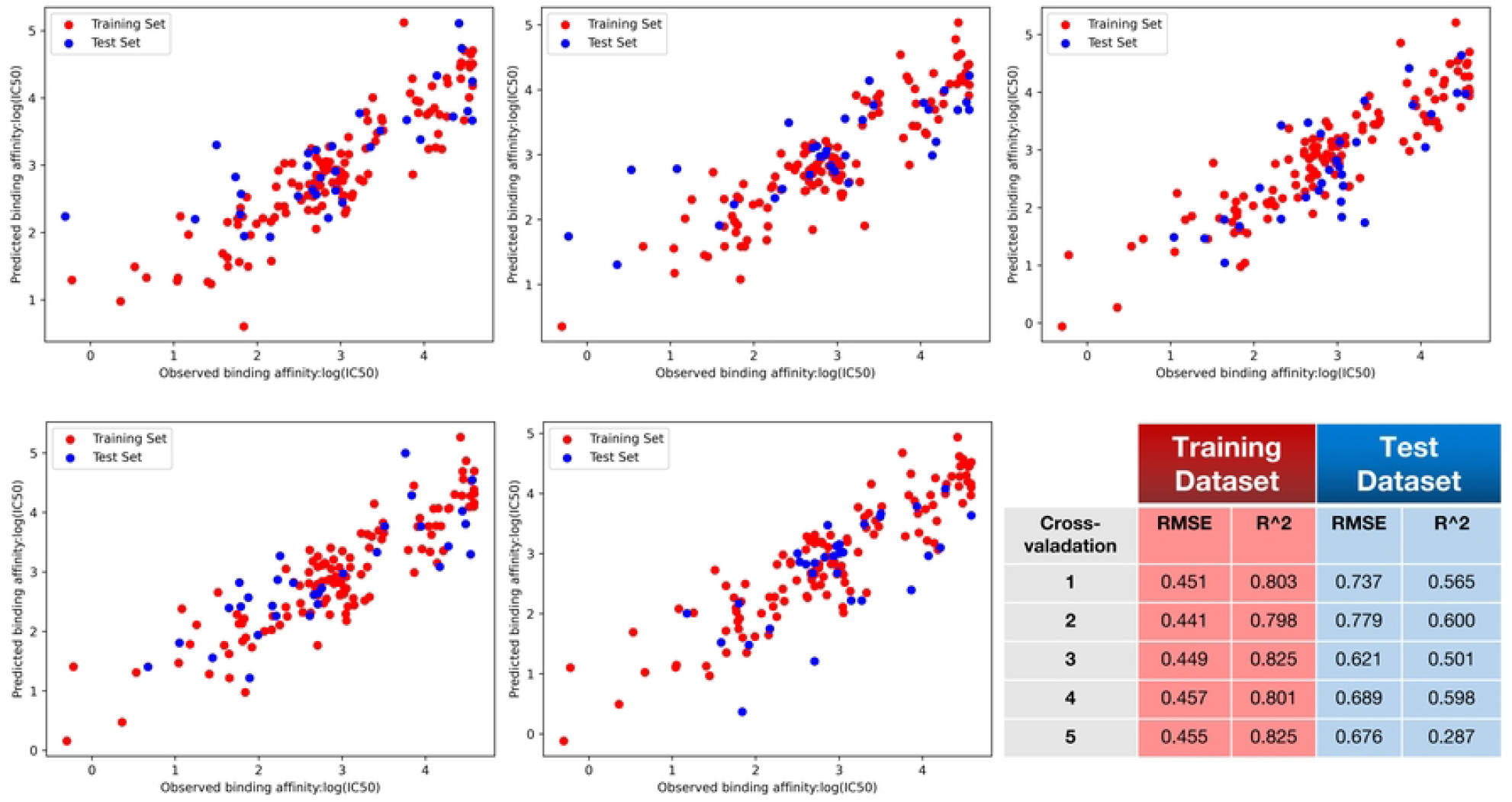
PLS model performance (5-fold cross validation) shown as actual vs. predicted log_10_(IC50). After splitting the model into 5 equal sized training and testing data sets, the correlation between predicted and experimentally determined IC50 values was calculated. Training dataset shown in red, testing in blue.

To provide a comprehensive comparison, the R^2^ values of the test dataset from various regressors employed by the Lazy Predict tool are plotted in Figure 7. The R^2^ and RMSE results that are comparable to those of the PLS model are summarized in Table 3.

**Table 3:** Performance of each regressor tested.

| Regressor | R-squared | RMSE |
| --- | --- | --- |
| SVR | 0.504 | 0.614 |
| Random Forest | 0.506 | 0.631 |
| Extra Trees | 0.515 | 0.608 |
| AdaBoost | 0.560 | 0.578 |

**Figure 7:**
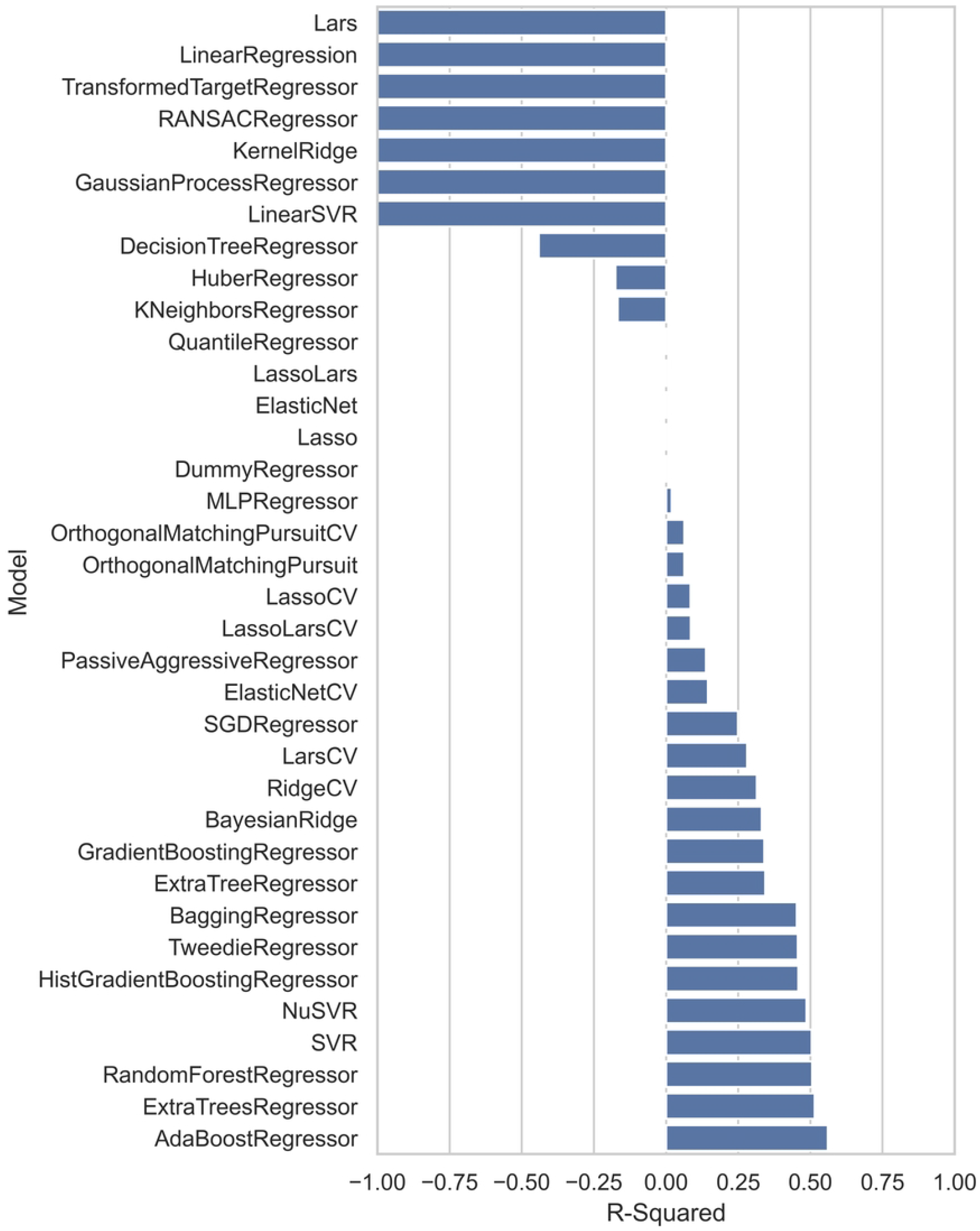
Model performance across various regressors, comparing R^2^ values for different algorithms tested on the same dataset.

## Discussion

Compared with other sequence-based prediction tools such as NetMHCPan [10,11], the most important improvement our model achieves is its ability to significantly expand the coverage of amino acid species in the involved peptides. Not only does it include the 20 canonical amino acids, but it also takes NCAAs into account without compromising structural accuracy. As long as the structure of an NCAA is known, applying this protocol to predict affinity is straightforward. Additionally, to make the model even more user-friendly, we have eliminated the need for MHC involvement in the model-building process. This means that, when compared with structure- or model-docking based approaches such as Rosetta FlexPepDock [15], our model provides results much faster with minimal human intervention. This is because our method does not require the provision and fine-tuning of large and complex protein structures, thereby accelerating the prediction process and reducing the potential for user error.

Despite these promising results, it is important to acknowledge that the current size of the training and test datasets is relatively small, which may limit the model’s performance. Although the Immune Epitope Database (IEDB) contains a substantial amount of data regarding peptide-MHC binding affinities, only a small percentage of these data includes quantitative binding values, and an even smaller portion pertains to peptides containing NCAAs. Collecting more data would enhance the model’s ability to capture a broader range of patterns and interactions, thereby improving its robustness and reliability.

Another future effort involves expanding the scope of the model to include MHCs from other species. For this study, we used data related solely to HLA-A0201 to ensure consistency, but extending the protocol to incorporate other MHC types would significantly widen the prediction coverage and improve the model’s reliability. By encompassing a larger variety of MHC alleles, we can better understand the nuances of peptide-MHC interactions across different biological contexts, making the model more universally applicable.

In conclusion, our model presents a notable advancement in peptide-MHC binding affinity predictions by expanding amino acid coverage and simplifying the prediction process. Future enhancements through increased dataset size and broader MHC coverage will further solidify its utility and accuracy, making it a powerful tool for computational immunology and related fields.

## Code Availability

The codes and corresponding data are available at https://github.com/meilerlab/ML_PLS_MHC_peptide_binding_pred with GNU General Public Licence Version 3.

## Acknowledgement

J.M. is supported by a Humboldt Professorship of the Alexander von Humboldt Foundation. J.M. acknowledges funding by the Deutsche Forschungsgemeinschaft (DFG) through SFB1423 (421152132), SFB 1664 (514901783), TRR (514664767), and SPP 2363 (460865652). J.M. is supported by the Federal Ministry of Education and Research (BMBF) through the Center for Scalable Data Analytics and Artificial Intelligence (ScaDS.AI), through the German Network for Bioinformatics Infrastructure (de.NBI), and through the German Academic Exchange Service (DAAD) via the School of Embedded Composite AI (SECAI 15766814). Work in the Meiler laboratory is further supported through the National Institute of Health (NIH) through R01 HL122010, R01 DA046138, R01 AG068623, U01 AI150739, R01 CA227833, R01 LM013434, S10 OD016216, S10 OD020154, S10 OD032234, 5T32HL144446-05. DGH is supported by NIH grants IH R35HL140016 and NIH AG076785, Z.S. thanks for the support of the Vanderbilt Data Science Postdoctoral Fellowship.

## Author contributions

Z.S., J.M. and DGH conceptualized and designed the study. S.J. and N.B. prepared the dataset. Z.S. and S.J. contributed to software development, model training, and analysis. S.J., Z.S., N.B., Y.L., C.M.. DGH and J.M. contributed to the writing and review of the manuscript. J.M. and DGH provided funding support

## References

1. Dendrou CA, Petersen J, Rossjohn J, Fugger L. HLA variation and disease. Nat Rev Immunol. 2018;18: 325–339. doi:10.1038/nri.2017.143

2. Charles A Janeway J, Travers P, Walport M, Shlomchik MJ. The major histocompatibility complex and its functions. Immunobiology: The Immune System in Health and Disease 5th edition. Garland Science; 2001. Available: https://www.ncbi.nlm.nih.gov/books/NBK27156/

3. Matsumura M, Fremont DH, Peterson PA, Wilson IA. Emerging principles for the recognition of peptide antigens by MHC class I molecules. Science (New York, NY). 1992;257: 927–934. doi:10.1126/SCIENCE.1323878

4. Archbold JK, Macdonald WA, Gras S, Ely LK, Miles JJ, Bell MJ, et al. Natural micropolymorphism in human leukocyte antigens provides a basis for genetic control of antigen recognition. J Exp Med. 2009;206: 209–219. doi:10.1084/jem.20082136

5. James EA, Rieck M, Pieper J, Gebe JA, Yue BB, Tatum M, et al. Citrulline-specific Th1 cells are increased in rheumatoid arthritis and their frequency is influenced by disease duration and therapy. Arthritis Rheumatol. 2014;66: 1712–1722. doi:10.1002/art.38637

6. Annet K, v F, AP de F, R L, CL G, J W, et al. DC isoketal-modified proteins activate T cells and promote hypertension. J Clin Invest. 2014;124: 4642–4656. doi:10.1172/JCI74084

7. Kacen A, Javitt A, Kramer MP, Morgenstern D, Tsaban T, Shmueli MD, et al. Posttranslational modifications reshape the antigenic landscape of the MHC I immunopeptidome in tumors. Nat Biotechnol. 2022; 1–13. doi:10.1038/s41587-022-01464-2

8. W Z, X S. Systematically benchmarking peptide-MHC binding predictors: From synthetic to naturally processed epitopes. PLoS computational biology. 2018;14. doi:10.1371/JOURNAL.PCBI.1006457

9. Mei S, Li F, Leier A, Marquez-Lago TT, Giam K, Croft NP, et al. A comprehensive review and performance evaluation of bioinformatics tools for HLA class I peptide-binding prediction. Brief Bioinform. 2020;21: 1119–1135. doi:10.1093/bib/bbz051

10. Hoof I, Peters B, Sidney J, Pedersen LE, Sette A, Lund O, et al. NetMHCpan, a method for MHC class I binding prediction beyond humans. Immunogenetics. 2009;61: 1–13. doi:10.1007/S00251-008-0341-Z

11. Nielsen M, Andreatta M. NetMHCpan-3.0; improved prediction of binding to MHC class I molecules integrating information from multiple receptor and peptide length datasets. Genome Medicine. 2016;8: 33. doi:10.1186/s13073-016-0288-x

12. Reynisson B, Alvarez B, Paul S, Peters B, Nielsen M. NetMHCpan-4.1 and NetMHCIIpan-4.0: improved predictions of MHC antigen presentation by concurrent motif deconvolution and integration of MS MHC eluted ligand data. Nucleic Acids Research. 2020;48: W449–W454. doi:10.1093/nar/gkaa379

13. O’Donnell TJ, Rubinsteyn A, Bonsack M, Riemer AB, Laserson U, Hammerbacher J. MHCflurry: Open-Source Class I MHC Binding Affinity Prediction. Cell Syst. 2018;7: 129–132.e4. doi:10.1016/j.cels.2018.05.014

14. Bloodworth N, Barbaro NR, Moretti R, Harrison DG, Meiler J. Rosetta FlexPepDock to predict peptide-MHC binding: An approach for non-canonical amino acids. PLOS ONE. 2022;17: e0275759. doi:10.1371/journal.pone.0275759

15. Raveh B, London N, Zimmerman L, Schueler-Furman O. Rosetta FlexPepDock ab-initio: Simultaneous Folding, Docking and Refinement of Peptides onto Their Receptors. PLOS ONE. 2011;6: e18934. doi:10.1371/journal.pone.0018934

16. Tengfei L, X P, L C, W T, S Q, L Y, et al. Subangstrom accuracy in pHLA-I modeling by Rosetta FlexPepDock refinement protocol. J Chem Inf Model. 2014;54: 2233–2242. doi:10.1021/CI500393H

17. Ferreira LLG, Andricopulo AD. ADMET modeling approaches in drug discovery. Drug Discov Today. 2019;24: 1157–1165. doi:10.1016/j.drudis.2019.03.015

18. Jiang L, Yu H, Li J, Tang J, Guo Y, Guo F. Predicting MHC class I binder: existing approaches and a novel recurrent neural network solution. Brief Bioinform. 2021;22: bbab216. doi:10.1093/bib/bbab216

19. Jing X, Dong Q, Hong D, Lu R. Amino Acid Encoding Methods for Protein Sequences: A Comprehensive Review and Assessment. IEEE/ACM Transactions on Computational Biology and Bioinformatics. 2020;17: 1918–1931. doi:10.1109/TCBB.2019.2911677

